# Comparative genomics studies provide insights into the taxonomic classification and secondary metabolic potential of five bioactive *Streptomyces* species isolated from the North-Western Himalaya

**DOI:** 10.1101/2024.05.28.596145

**Authors:** Aasif Majeed Bhat, Mariam Azeezuddin Haneen, Aehtesham Hussain, Gaurav Sharma, Qazi Parvaiz Hassan

## Abstract

The linear genome of genus *Streptomyces* members has the potential to encode diverse and novel biosynthetic gene clusters of invaluable antimicrobial and therapeutic significance. The use of limited taxonomic markers makes the precise identification of these miracle microbes very challenging. In the ongoing omics era, genome sequencing and *in-silico* analysis of these potential antibiotic producers provide deeper insights into their taxonomy, functional capabilities, and potential for antibiotic production. Here this study presents a multifaceted approach for proper taxonomic identification and genomic and bioinformatic analysis of five bioactive *Streptomyces* species collected from different sampling sites in the high-altitude oligotrophic North-Western Himalaya, Kashmir, India. We used polyphasic taxonomic classification approaches, such as phylogenetic markers (16S rDNA and gyrase B), average nucleotide identity (ANI) estimation, and digital DNA-DNA hybridization (dDDH), which revealed accurate taxonomic placement of five *Streptomyces* species, named as, *Streptomyces violarus* ASQP_29, *S. rhizosphaerihabitans* ASQP_78, *S. fulvoviolaceus* ASQP_80, *S. mirabilis* ASQP_98, and *S. thajiwasiensis* ASQP_92. Amongst these, one notable finding is the discovery of a novel species proposed as *Streptomyces thajiwasiensis* sp. nov. ASQP_92. In addition, our study presents the first genome announcement report and analysis for *S. rhizosphaerihabitans* ASQP_78. Genomic annotation highlighted the presence of an exceptionally high number of poorly characterized genes and hypothetical proteins, indicating their potential for undiscovered biotechnological applications. Clusters of orthologous groups (COG) and gene ontology (GO) analysis provided insights into their varied functional roles in metabolism, signaling, information storage and secondary metabolite biosynthesis. Domain-based functional characterization further detailed their involvement in various biological processes particularly in antibiotic biosynthesis, transport, and resistance. Biosynthetic gene clusters (BGC) analysis demonstrated their diverse metabolite biosynthetic capabilities and identified both unique and conserved BGCs emphasizing the species-specific roles in bioactive metabolite production and the potential of orphan BGCs in novel drug discovery.

**Graphical abstract:**
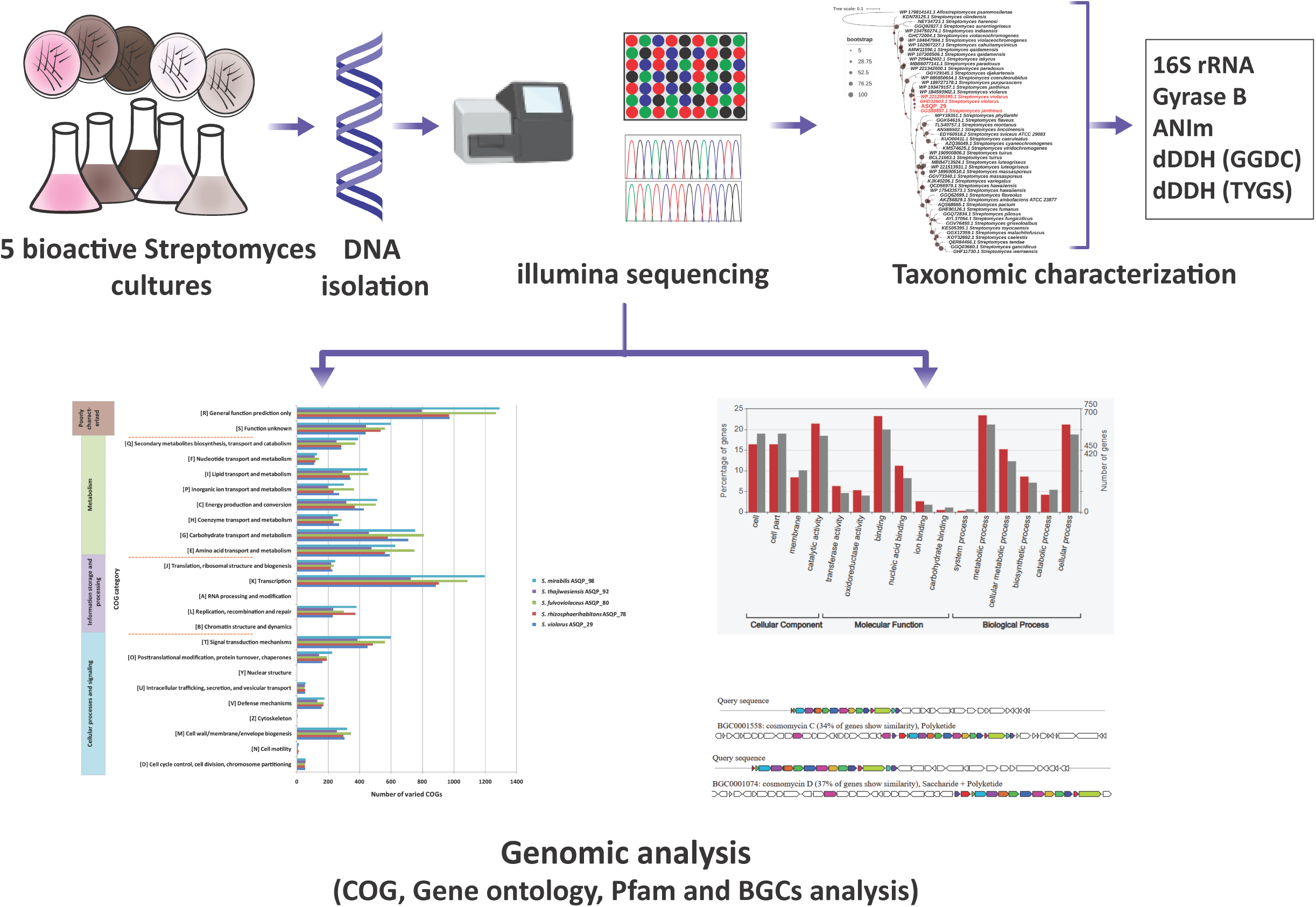
Genomic analysis and taxonomic affiliations of five bioactive *Streptomyces* species isolated from high altitudes of the North Western Himalaya

## Introduction

Genus *Streptomyces* members are gram-positive aerobic bacteria of the phylum Actinomycetota (earlier known as phylum Actinobacteria) (Oren & Garrity, 2021; Panda et al., 2022) having filamentous linear and branched mycelial morphology and aerial hyphae that bear conidial spores at maturity (Witt & Stackebrandt, 1990). The contrasting peculiar characteristic of *Streptomyces* is the high GC percentage, and linear chromosome structure with unique terminal-inverted repeats at both ends of the chromosome (Lin et al., 1993; Qazi et al., 2019). These linear structures encode secondary metabolite biosynthetic gene clusters (SMBGC) that are of great pharmacological and commercial importance for the production of different classes of antibiotics, antiparasitics, herbicides, pharmacologically important bioactive metabolites, and enzymes valuable in food and other industries (Demain, 1999). Therapeutically invaluable *Streptomyces* species encode enzyme components and catalytic domains of BGCs notably biosynthetic modular PKS, NRPS, and hybrid PKS-NRPS systems which have shown a high degree of mechanistic, structural, and functional similarity to natural products for pathogen inhibition (Hill, 2006; Lee et al., 2020; McDaniel et al., 2005; Weissman & Leadlay, 2005). Recent advances in genome sequencing technologies and *in-silico* genome analysis have revealed that *Streptomyces* genomes contain numerous silent small metabolite biosynthetic gene clusters (smBGCs) that remain uncharacterized under conventional laboratory conditions (Lee et al., 2020). *Streptomyces* genomic study underpins the importance of cryptic BGCs and the modular structural prediction and domain characterization of these gene clusters (Jenke-Kodama & Dittmann, 2009). This in-depth understanding of biosynthetic systems encoded in these megasynthase clusters, along with the specialized development of bioinformatics tools, has paved the way for the possible use of genomic sequences in the development of new metabolites. Both prospective and retrospective approaches to genome mining can be employed for natural product exploitation in cultured and uncultured *Streptomyces*. However, the preferred strategy for cultured *Streptomyces* involves retrospective genome mining, wherein high-throughput and/or flask-based growth and fermentation of cultured *Streptomyces* guide screening for the evaluation of their natural product potential against various test pathogens (Foulston, 2019).The genomes of selected bioactive strains are then sequenced, and the nearly full genomic and biosynthetic gene cluster (BGC) characteristics of an organism are evaluated with a focus on nonribosomal peptide synthetases (NPRS), polyketide synthases (PKS), and NRPS-PKS hybrids, among others (Ward & Allenby, 2018). This gives a complete understanding of the expressed and unexpressed natural products potential of *Streptomyces*. Further, the golden era of drug discovery had accelerated the pace with which the *Streptomyces* organisms were isolated. However, the limited taxonomic parameters of that time made the identification of these miracle microbes challenging (Law et al., 2018). This bolstered the concept of phylogeny-based genome mining or evomining with taxogenomic classification, where methods such as ANI calculation and *in-silico* DNA-DNA hybridization aid in the accurate identification of isolated *Streptomyces* and the detection of novel taxa. The use of whole-genome sequencing (WGS) has revolutionized the field of taxonomy, particularly for the phylum Actinobacteria, and has provided a more reliable means of identifying and describing new species (Kusuma et al., 2021). While the traditional approach of using 16S rRNA gene sequences and phylogeny is still useful, it is often inadequate for differentiating between closely related species within the genus *Streptomyces* (Kämpfer & Labeda, 2006). The taxogenomic approach using WGS provides a more comprehensive and accurate means of characterizing bacterial species and has been used to support other polyphasic taxonomic data (Kusuma et al., 2021). The use of bioinformatic tools and WGS analysis has not only benefited the field of taxonomy but has also contributed to our understanding of developmental and evolutionary processes, as well as the ecology, physiology, and metabolite production of *Streptomyces* (Otani et al., 2022). With the aid of WGS, researchers can obtain detailed information on the genetic makeup of bacteria, which can provide insights into their functional capabilities, potential for antibiotic production, and adaptation to environmental conditions. This knowledge can be used to guide the development of new antibiotics and other biologically active compounds, while also providing insights into the evolutionary relationships and ecological roles of *Streptomyces* (Otani et al., 2022).

In our recent work (Bhat et al., 2024), we have performed the biochemical characterization of these five *Streptomyces* strains and several other isolates. This study has also identified their minimum inhibitory concentrations (MICs) against test strains of microorganisms by performing *in vitro* antimicrobial activity. To further continue our exploration, the present study entails these five *Streptomyces* for genomic analysis because; 1) these *Streptomyces* are isolated from high altitude, oligotrophic and under-explored extreme terrestrial habitats of North Western Himalaya (NWH); 2) these are bioactive isolates showing antimicrobial potential against a panel of gram-positive and gram-negative pathogens; 3) several bio evaluation studies by our research group has shown that *Streptomyces* from NWH produces bioactive natural products of immense biotechnological and pharmacological potential and isolates from these high altitude regions secrete secondary metabolites that exhibit significant antimicrobial, anti-TB and antiproliferative activities (Hussain et al., 2017; Hussain, Dar, et al., 2019; Hussain, Rather, et al., 2019; Shah et al., 2016; Wani et al., 2019). But little has been done to unsnarl the genomic complexity and resolve molecular-based multifunctional consortia of these high-altitude biosynthetic producers; 4) further *Streptomyces* genomes have a rich history of producing FDA-approved antibiotics(Moumbock et al., 2021)and hence the genomic investigations into these potential producers is a step forward in elucidating the biosynthetic capacity of these antimicrobial-producing microorganisms and 5)The comparative genomic analysis of bioactive *Streptomyces* from NWH will also help to provide genomic insights into the potential producers of pharmacologically important metabolites and this will go along in understanding the application of these microbes in drug discovery paradigms (Bhat et al., 2022). Overall, following our previous experimental work (Bhat et al., 2024), this study focuses on taxonomic classification, genome sequencing, annotation, and secondary metabolite clusters identification of five *Streptomyces* spp. to explore potential novel future metabolites.

## Methodology

### Sample isolation, bacterial culturing, and organism identification

Soil samples were collected from oligotrophic sites at high altitudes (2128-4390 m) (Bhat et al., 2024)in the North Western Himalayas at depths of 5-20 cm and were then serially diluted as 5-7 dilutions each down to 10^-1^. The dilutions were spread plated on actinobacteria-specific media (Bhat et al., 2024), and culture plates were inspected every two days for growth of actinobacteria colonies for 6-20 days at 28°C. Putative actinobacteria colonies on culture plates were selected, subcultured and purified. *Streptomyces* were identified based on their morphology, including the transition from the substrate to aerial mycelial patterns, sporulation in aerial mycelia, pigmentation, and observations from Gram staining, meticulously examined under a binocular Leica microscope model DM 5500 (Bhat et al., 2024). The selective isolates of *Streptomyces,* as mentioned in the previous research article were isolated from different sampling sites as ASQP_29 from S1 (Sinthan top); ASQP_78, ASQP_80, and ASQP_98 from S4 (Peer Ki Gali) and ASQP_92 from S2 (Thajiwas glacier) (Bhat et al., 2024). The selective isolates of *Streptomyces* were preserved as 2ml aliquots in 20% glycerol/PBS (v/v) at -80°C.Whole genomic DNA was extracted from the pure cultures using DNeasy Power Soil Pro Kit, Qiagen. DNA was quantified (NanoDrop Spectrophotometer ND-1000, Thermo Scientific), and run on 0.8% gel to check for purity and to verify the high molecular weight. Using a universal primer set of 27F (5′-AGAGTTTGATCCTGGCTCAG-3′) and 1492R (5′-GGTTACCTTGTTACGACT T-3′), nearly full-length 16S rRNA non-coding DNA was amplified at in-house Sanger sequencing facility at NCMR-NCCS, India. 25 µL PCR reaction was performed in 96 well Thermal cycler (Applied Biosystems) with 1 µL DNA template (<50 ng µL−1), 2.5 µL Taq polymerase assay buffer, 1µL each 10 µM primers, 2 µL dNTPs, 1.5 µL MgCl, 0.25 µL Taq DNA Polymerase (QIAGEN) and 15.75 µL nuclease-free water. Amplification profile with initial denaturation at 94 °C (3mins), followed by 30 cycles of denaturation at 94 °C (30s), annealing at 55 °C (30s), extension at 72 °C (90s), and final extension at 72 °C (5min) followed by infinite hold at 10°C. Amplicons were confirmed on 1% agarose electrophoresis gel before cleanup by the PEG/NaCl purification system. The resultant sequences (>1400nt) were searched for homology against different strains using the NCBI nucleotide blast search database.

### Genomic sequencing and assembly validation

Two different Illumina sequencing platforms were employed for whole genomic sequencing of selected five bioactive *Streptomyces* species, Illumina HiSeq for ASQP_29 and ASQP_80 and Illumina MiSeq for ASQP_78, ASQP_92, and ASQP_98. 100ng of intact quality DNA was sheared enzymatically by NEBNext Ultra II kit to approximately 200-300 bp size. The end-repair 3‘-5‘ exonuclease activity removes 3‘overhangs and polymerase activity fills in 5‘ overhangs. The resultant blunt-ended fragments undergo adenylation at the 3‘ end, loop adapters are ligated and cleaved with uracil-specific excision reagent-USER enzyme. Samples are purified using AMPure beads. To enrich the DNA, six cycles of PCR reaction were performed using Illumina universal primer, NEBNext Ultra II Q5 master mix, and sample-specific octamer primers. The amplified DNA library was further cleaned by AM pure beads and eluted in 15 µLs of 0.1X TE buffer. To check the library QC, 1µl of DNA was quantified using QUBIT 3 Fluorometer with dS DNA HS reagent and fragments were analyzed on Agilent 2100 Bioanalyzer using Agilent DNA 7500 chip.

Sequencing read counts, read length, and coverage were estimated followed by the quality trimming and adapter removal of raw reads using Trimmomatic-0.39v (Bolger et al., 2014) and quality reassessment of the reads using FASTQC (Andrews, 2010). *De novo* assembly was constructed using the Unicycler assembler tool (Wick et al., 2017) and assembly validation was performed using QUAST-5.0.2v (Gurevich et al., 2013). To predict the identification of probable genetic elements like CDS (protein-coding genes), RNAs, repeats, etc., the rearranged contigs FASTA assemblies of all five strains were submitted to the RASTtk server (Aziz et al., 2008) for annotation of the genomes. The resultant RAST gene and protein files were subjected to different tools and databases as mentioned below for further functional characterization.

### 16S rRNA and gyrase B protein based phylogeny

16S rRNA sequences and DNA gyrase B protein sequences of all the five *Streptomyces* strains were extracted from RAST annotation files (RNA FASTA and amino acid FASTA). BLASTn and BLASTp were respectively performed against NCBI’s 16S rRNA and NR databases. Uncultured and unclassified hits were unchecked, redundant hits were removed, and unique top 50 hits were selected and downloaded. *Allostreptomyces psammosilenae* (WP 179814141.1) was taken as an outgroup for this study. All extracted sequences from *Streptomyces* strains and outgroup were subjected to multiple sequence alignment (MSA) using MUSCLE-v3.8.1551 with all default parameters (maxiters default value: 16). Model selection for each MSA was done using MEGAX software (Tamura et al., 2021). An optimum model was selected for the phylogenetic construction of each *Streptomyces* species and tree inference was done using RaxML (Stamatakis, 2015) with an optimized model and 100 bootstrap values followed by visualization in iTOL (Letunic & Bork, 2021).

### Species delineation using Maximal Unique Matches (MUMmer), Genome-to-Genome Distance Calculator (GGDC), and Type Strain Genome Server (TYGS)

An in-house constructed script and prokaryotes.txt file (at NCBI FTP site) were employed to download all reported *Streptomyces* genomes and a python-based tool pyani v0.3 (Pritchard et al., 2016) for the MUMmer module was run to calculate Average Nucleotide Identity-ANIm values and estimate the similarity percentage of query strains vis-a-vis other *Streptomyces* (Delcher et al., 2003). Subsequently, the query strains were checked for *in-silico* DNA-DNA hybridization (DDH) using type strain-dependent TYGS or type strain independent GGDC web servers (Meier-Kolthoff & Göker, 2019).

### COG (Clusters of Orthologous Groups) analysis

Protein sequences of all five strains were searched for COG annotations using RPS-BLAST+ with e-value 1e^-5^against NCBI’s CDD database (COG profiles) (My et al., 2021). For each protein, PSSM-IDs were mapped to the cddid.tbl file to extract corresponding COG categories. These categories were then mapped to the whog file, allowing for the elucidation of orthologous functional categories. The number of proteins belonging to each functional category was calculated and assigned to the main categories. Bar plots were plotted using the ggplot2 package in R (v4.0.3).

### GO (Gene Ontology) analysis

Protein sequences for each organism were subjected to HMMER2GO (v0.18.0) (Evan, 2022; Prakash et al., 2017) and respective Gene Ontology-GO IDs were assigned to all the coding sequences of five *Streptomyces* strains. Using the WEGO web server (Ye et al., 2018) GO IDs are attributed to corresponding GO terms and the number of genes belonging to each GO term are calculated. Representative bar plots were drawn using the ggplot2 package in R (v4.0.3).

### Pfam (Protein families) domain analysis

*For in-silico* protein annotation and identification of different protein families, the protein sequences of five *Streptomyces* strains were subjected to the hmmscan program in the HMMER suite (Prakash et al., 2017) against Pfam-A v 35.0 databases with a threshold E-value of 1e^-5^ (Mistry et al., 2021). The identification and distribution of different protein families were analyzed.

### BGC (Biosynthetic Gene Cluster) analysis

The bioactive secondary metabolite gene cluster prediction analysis was done by genome mining of *Streptomyces* strains against the MIBiG database using the antiSMASH (antibiotics and secondary metabolites analysis shell) tool and different Biosynthetic Gene Clusters (BGCs) were categorized (Blin et al., 2013; Medema et al., 2011).

## Results

Our investigation commenced with the morphological analysis and microscopic assessment of five selected bioactive *Streptomyces* species. The identification of pigmented hyphal morphology branched mycelial formations, and spore-producing mycelia confirmed the similarity of these species structures to those typical of the *Streptomyces* genus (Fig. 1).

**Figure 1.**
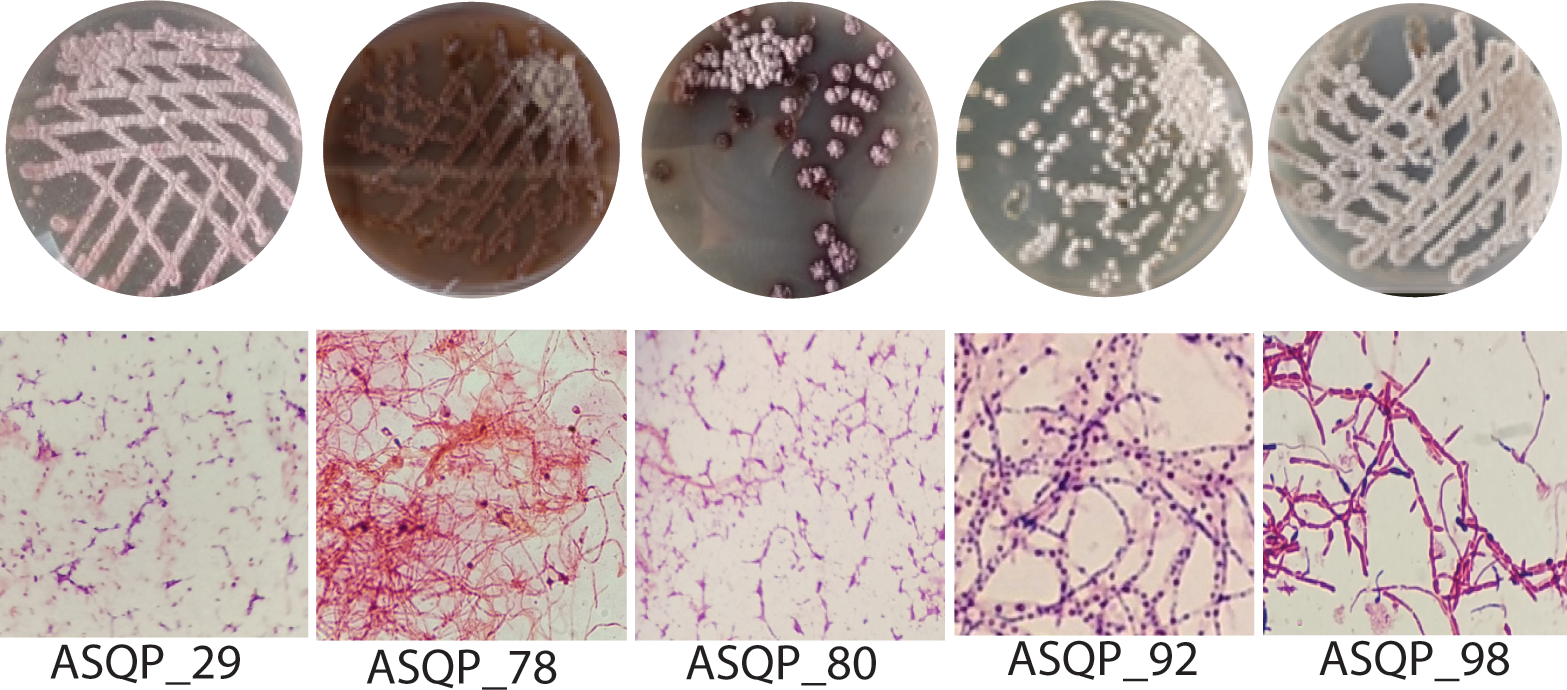
Morphological and microscopic features of five bioactive *Streptomyces* species.

### 16S rDNA and gyr-B based phylogenies of the five bioactive *Streptomyces* species accurately delineated their taxonomic classifications, assigning them to their respective neighbouring *Streptomyces* species

Phylogenetic analysis based on 16S rDNA showed that ASQP_29 (1338 nt) strain clustered together with *S. violarus* and *S. arenae* species. ASQP_29 strain exhibited 100% identity to each *S. violarus* (NR_114829.1), *S. arenae (*NR_025494.1), and *S. violarus* (NR_041116.1). Its percentage identity with the closely related *S. purpurascens* strains was 99.92% (Fig. 2a). ASQP_78 (1532 nt) strain clustered with *S. rhizosphaerihabitans (*NR_151948.1) with a percentage identity of 99.80%. The percentage identity with both *S. olivochromogenes* strains was 99.25% (Fig. 2b). ASQP_80 (1534 nt) strain formed a clade with *S. triticiradicis* (NR_169450.1) with 98.29% identity, however, the strain depicted the highest percentage identity with *S.pseudovenezuelae* (NR_041090.1) (99.39%), followed by *S.pseudovenezuelae* (NR_114832.1) (99.38%), *S.novaecaesareae* (NR_041124.1) (99.2%), *S.canus* (NR_043347.1) (99.1%), *S.canus* NR_12259.1 (99.1%), *S.canus* (NR_041085.1) (99.1%) and *S. resistomycificus* (NR_112287.1) (99.0%) (Fig.2c).

**Figure 2.**
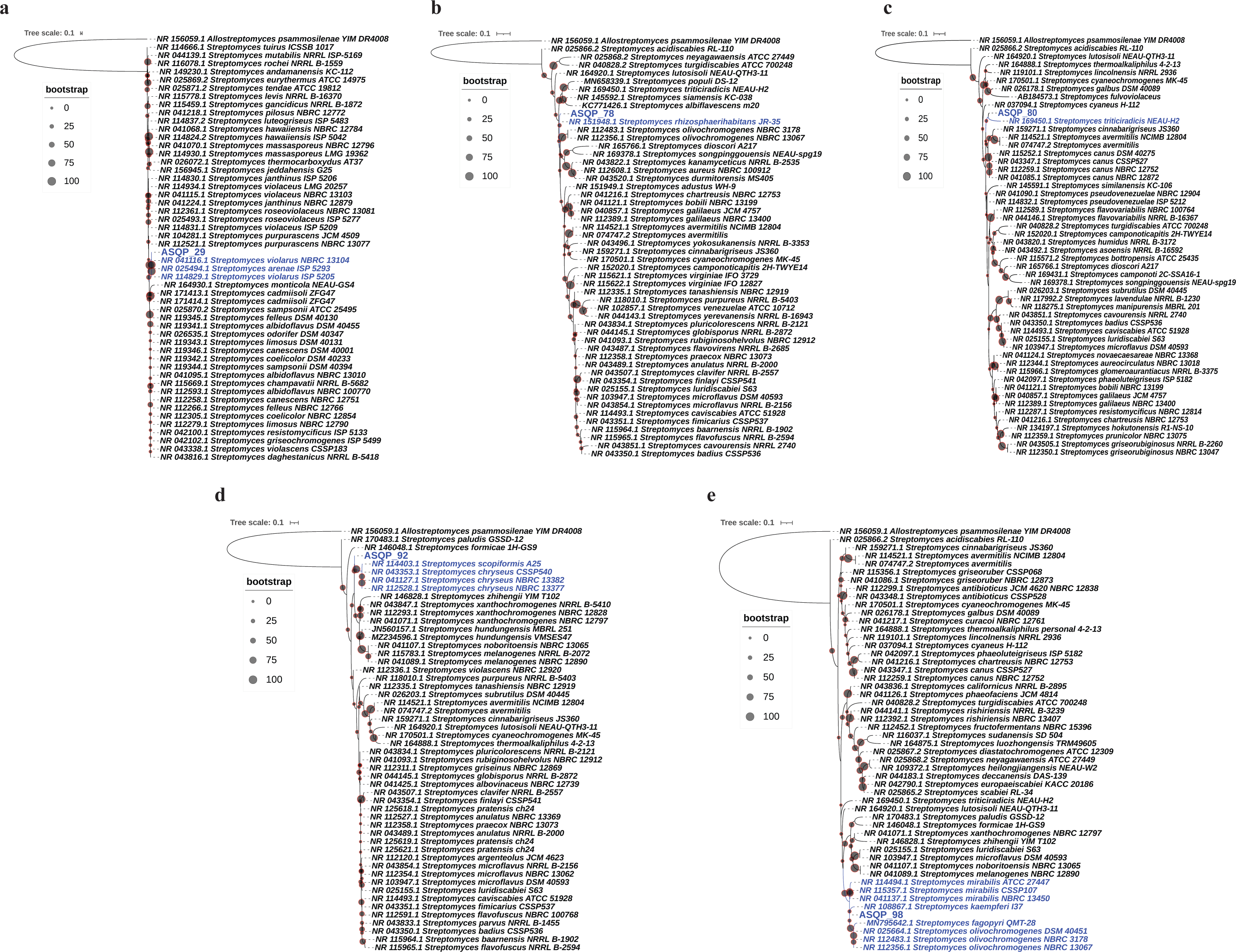
16S rRNA based phylogeny for ASQP_29 (a), ASQP_78 (b), ASQP_80 (c), ASQP_92 (d) and ASQP_98 (e). Maximum Likelihood method was used to compute evolutionary distances and the final evolutionary analysis was inferred using RAxML with 100 bootstrap replicates. The tree was visualized, manipulated, and annotated in the Interactive Tree of Life (iTOL). Colored nodes represent strains that form the closest possible clade/s with the query *Streptomyces* species. *Allostreptomyces psammosilenae* serves as the outgroup species.

ASQP_92 (1532 nt) strain clustered with *S. scopiformis* and *S. chryseus* showcasing 98.9% identity with both the strains, followed by 99.46% identity with other close relatives, i.e., *S. xanthochromogenes* (NR_041071.1), *S. xanthochromogenes* (NR_043847.1) and *S. xanthochromogenes* (NR_112293.1) (Fig. 2d). ASQP_98 (1347 nt) clustered with *S. kaempferi, S. fagopyri* QMT-28, and three different strains of each *S. mirabilis* and *S. olivochromogenes*. The strain formed the highest percentage identity with *S. mirabilis* NR_115357.1 (99.9%), *S. mirabilis* NR_041137.1 (99.9%), *S. mirabilis* (NR_114494.1) (99.8%), followed by *S. lutosisoli* (NR_164920.1) (99.6%), and *S. kaempferi* (NR_108867.1) (99.5%) (Fig. 2e).

### Phylogeny designation based on the B subunit of the DNA gyrase protein (gyrB) identity

Along with 16S-based phylogeny, which relies on DNA sequences of the query strains for *Streptomyces* species identification, we also utilized the amino acid sequences of the gyrB gene as a phylogenetic marker due to redundancy in the genetic code. Strain ASQP_29 (694 aa) clustered closely into three separate clades with *S. janthinus, S. violarus* and *S. purpurascens,* however the closest clade formation was observed with *S. janthinus* and *S. violarus*. The strain exhibited highest identity with *S. violarus* GHD_32603.1 (99.71%), followed by *S. janthinus* GGS_58857.1 (99.57%), *S. violarus* WP_221299195.1 (99.57%), *S. violarus* WP_184593902.1 (99.56%), *S. Janthinus* WP_193479157.1 (99.4%) and *S. purpurascens* WP_189727178.1 (98.1%) (Fig. 3a). Strain ASQP_78 (687 aa) formed a clade with *S. avermitilis* spp. but exhibited highest percentage identity with *S. triticiradicis* KAB_1986649.1 (96.51%) followed by *S. aureus* WP_037614948.1 (96.22%) and *S. populi* PKT_69761.1 (96.22%); however, the percentage identity with *S. avermitilis* KUN_52886.1 was 95.29% and with *S. avermitilis* WP_010985746.1 was 95.09% (Fig. 3b). Strain ASQP_80 (687 aa) formed a separate clade with *S. fulvoviolaceus* WP_106981039.1 (percentage identity 99.85%). The percentage identity with all other strains was less than 98% (Fig. 3c). Strain ASQP_92 (687 aa) clustered closely into three separate clades with *S. hundungensis*, *S. xanthochromogenes* and *S. violascens,* however, the closest clade formation was observed with *S. hundungensis* spp. The percentage identity was highest for *S. hundungensis* AYG_81752.1 (99.85%), *S. hundungensis*WP_174248584.1 (99.71%) followed by *S. xanthochromogenes* GGY_51042.1 (98.11%), *S. violascens* GGU_42021.2 (96.94%) and *S. violascens* WP_189969864.1 (96.76%). The percentage identity with all other strains was less than 96% (Fig. 3d). Strain ASQP_98 (694 aa) formed separate close clades with *S. mirabilis* and *S. olivochromogenes,* however, it is forming a closest clade with *S. mirabilis*. The percentage identity was highest and identical for *S. mirabilis* GHD_76724.1 and *S. olivochromogenes* KUN_43084.1 (99.57%for both strains). The percentage identity for *S. mirabilis* WP_099925533.1 strain was 99.42%, however, the identity percentage for all other strains was less than 96.5% (Fig. 3e).

**Figure 3.**
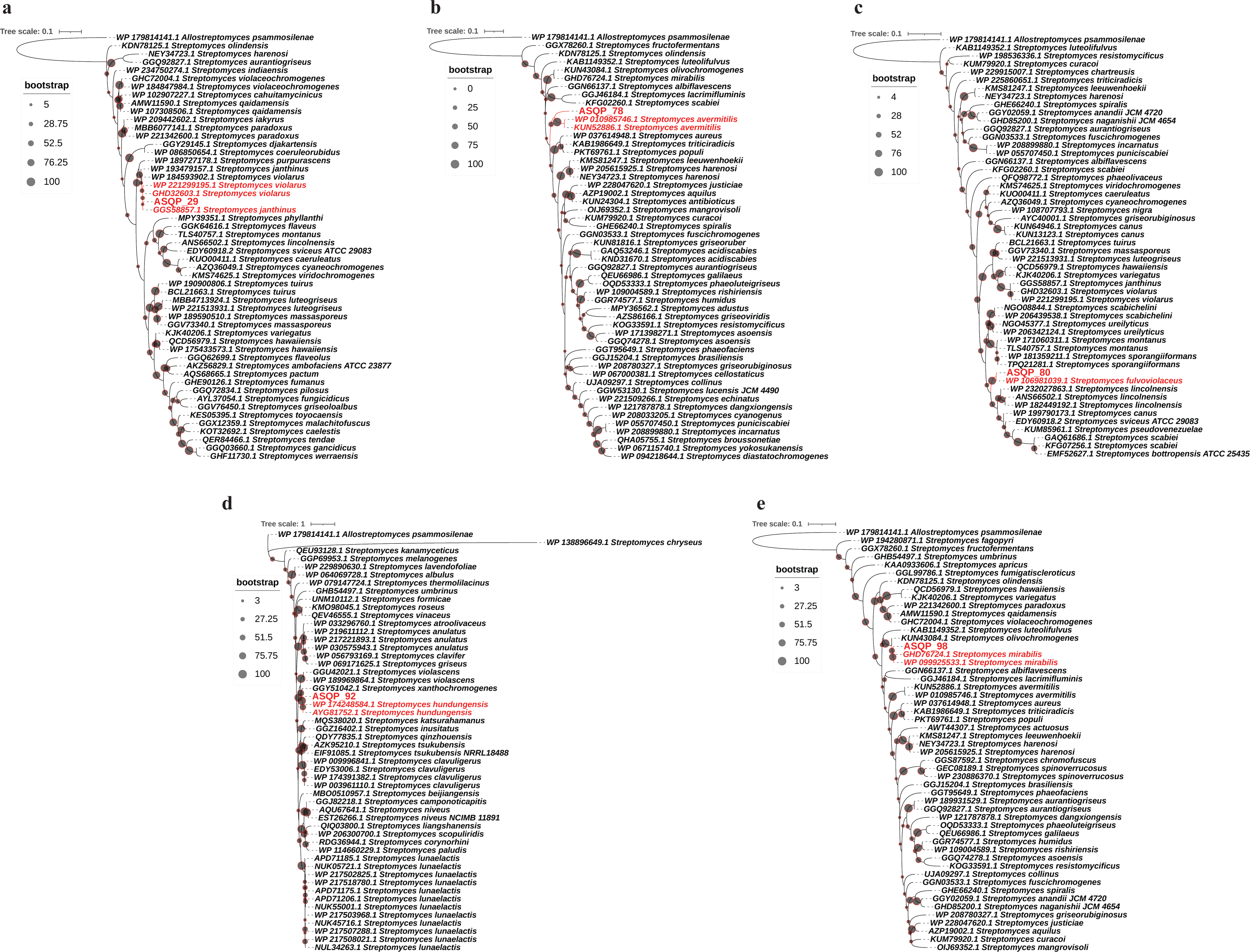
Gyrase B protein-based phylogeny for ASQP_29 (a), ASQP_78 (b), ASQP_80 (c), ASQP_92 (d) and ASQP_98 (e). Maximum Likelihood method was used to compute evolutionary distances and the final evolutionary analysis was inferred using RAxML with 100 bootstrap replicates. The tree was visualized, manipulated, and annotated in the Interactive Tree of Life (iTOL). Colored nodes represent strains that form the closest possible clade/s with the query *Streptomyces* species. *Allostreptomyces psammosilenae* serves as the outgroup species.

**Figure 4.**
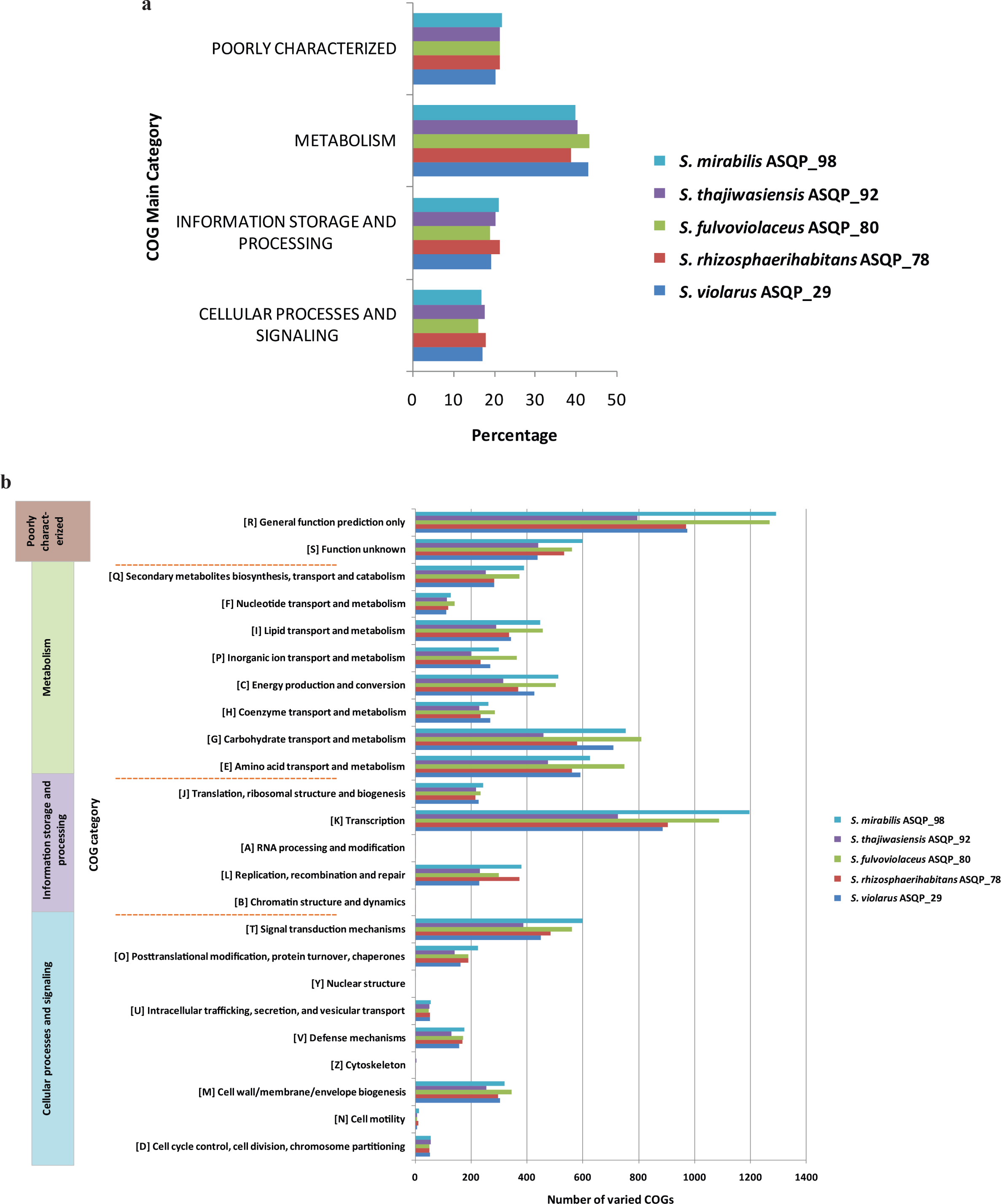
Comparative functional annotation using COG main and sub categories distribution among five *Streptomyces* species. a) Percentage of COGs assigned to different main categories. b) Number of varied COG subcategories shown as horizontal clustered bars along with their corresponding single letter codes in square brackets. The main categories are listed in colored boxes and are separated by red dotted lines.

### Average Nucleotide Identity (ANI) estimation evaluated the top closest relatives for each *Streptomyces* strain

The average nucleotide identity index values based on MUMmer (ANIm) were evaluated using an in-house pyani script and the top phylogenetically closest relatives for each were summed up in Supplementary Table 1. Strain ASQP_29 exhibited highest ANI value with *S. violarus* (83.1%), followed by *S. janthinus* (80.7%) and *S. purpurescens* (76.7%); strain ASQP_78 exhibited highest ANI value with *S. populi* (66.0%), followed by *S. triticiradicis* (65.1%) and *S. aureus* (62.8%); strain ASQP_80 exhibited highest ANI value for *S. fulvoviolaceus* (88.0%), followed by *S. lincolnensis* (60.5%), *S. resistomycificus* (59.9%) and *S. pseudovenezuelae* (58.9%); strain ASQP_92 exhibited highest ANI value with *S. hundungensis* (78.5%), followed by *S. xanthochromogenes* (61.7%) and *S. violascens* (58.8%); strain ASQP_98 exhibited highest ANI value with *S. mirabilis* (80.5%) followed by *S. olivochromogenes* (78.7%) and *S. fagopyri* (61.7%).

### GGDC estimated different digital DNA-DNA Hybridization (dDDH) relatedness characteristics for each *Streptomyces* species

*In-silico* based digital DNA-DNA hybridization (dDDH) values using DSMZ Genome to Genome Distance Calculator (GGDC v3.0) were evaluated and the recommended formula 2 [(identities/HSP (high-scoring segment pairs) length) which is preferred against the incomplete draft genome assemblies] were estimated and summed up in Supplementary Table 2. The strain ASQP_29 exhibited the dDDH values of 71.8% (C.I. model 68.8-74.6%) to *S. violarus*; 77.1% (C.I. model 74.1-79.8%) to *S. janthinus* and 62.7% (C.I. model 59.8-65.5%) to *S. purpurescens*. Strain ASQP_78 exhibited the dDDH values of 40.4% (C.I. model 37.9-43.0%) to *S. populi*; 40.4% (C.I. model 37.9-42.0%) to *S. triticiradicis*. The dDDH values and other formula 2 parameters for the rest of the strains against this query strain are very low. The strain ASQP_80 exhibited the highest dDDH values of 94.8% (C.I. model 93.2-96.1%) to *S. fulvoviolaceus*. The dDDH values for the rest of the strain were less than 32%. ASQP_92 exhibited less than 50% dDDH values to all the target strains and the highest dDDH value of 49.9% (C.I. model 47.2-52.5%) was reported to *S. hundungensis*. ASQP_98 exhibited the dDDH values of 68.1% (C.I. model 65.1-70.9%) to *S. mirabilis*; 68.1% (C.I. model 65.1-70.9%) to *S. olivochromogenes*; the dDDH values to rest of the strains were less than 34%.

### TYGS estimated pairwise digital DNA-DNA Hybridization (dDDH) values between query genomes and the top type stains genomes

*In-silico* pairwise digital DNA-DNA hybridization values using TYGS (TYGS v3.0) were further evaluated and all three formula results were estimated and summed up in Supplementary Table 3. The strain ASQP_29 exhibited the dDDH values for formula d_4_ of 77.1% (C.I. model 74.1-79.8%) to *S. janthinus*; 71.8% (C.I. model 68.8-74.6%) to *S. violarus* spp. and 62.7% (C.I. model 59.8-65.5%) to *S. purpurescens*. Thus *S. violarus* shows the most preferred Genome BLAST Distance Phylogeny (GBDP) value, owing to its comparable d_0_ andd_6_ formula values. Likewise for formula d_4_, the strain ASQP_78 exhibited the dDDH values of 40.4% (C.I. model 37.9-42.9%) to *S. triticiradicis*; 40.4% (C.I. model 37.9-43.0%) to *S. populi*. The dDDH values and other formula parameters for the rest of the strains against this query strain are lower than 33% and thus amount to incompetent phylogenomic comparison. The strain ASQP_80 exhibited the lowest dDDH values for all the reference type strains and the highest d_4_ value of 31.3% was reported for *S. resistomycificus, S. canus* and *S. ciscaucasicus*. The dDDH values for the rest of the strain were less than 31%. ASQP_92 exhibited less than 37 % d_4_ dDDH values to all the target strains and the highest dDDH value of 36.3% was reported to *S. violascens* and *S.michiganensis*. ASQP_98 exhibited the highest dDDH values of 68.1% (C.I. model 65.1-70.9%) to *S. mirabilis*; 68.0% (C.I. model 65.1-70.9%) to *S. olivochromogenes*; the dDDH values to the rest of the strains were less than 34%.

To summarize, the combination of phenotypic, phylogenetic, and taxogenomic analysis revealed that the strain ASQP_29 belonged to *S. violarus*, strain ASQP_78 to *S. rhizosphaerihabitans*, strain ASQP_80 to *S. fulvoviolaceus* and strain ASQP_98 to *S. mirabilis*; however, the strain ASQP_92 was different from all related strains under study and known ones. This strain could not be categorized within any of the known strains and therefore, represents a novel species within the genus *Streptomyces*. We propose the name *Streptomyces thajiwasiensis* sp. nov.ASQP_92, owing to its origin from the Thajiwas glacier in the Kashmir region of the Jammu and Kashmir state in India.

### Genome assembly information and annotation statistics of five *Streptomyces* species

The assessment of genome assembly information and annotation statistics are summed up in Table 1. All five *Streptomyces* strains exhibit a GC percentage greater than 70%, which is a typical feature of the genus *Streptomyces*. The genome of *S. violarus*ASQP_29 was assembled into ∼8.9 Mb within 585 contigs (N50 value of 27,723 and L50 value of 100) and further annotated into 8,898 genes and 61 RNAs. For *S. rhizosphaerihabitans* ASQP_78, the assembled genome was of ∼10.2 Mb within 244 contigs (N50 value of 147,426 bp and L50 value of 23) which were further annotated into 9,853 genes and 79 RNAs. *S. fulvoviolaceus* ASQP_80 genome was assembled into a draft genome of ∼11.0 Mb within 320 contigs (N50 value of 69,118 and L50 value of 51) followed by annotating a total of 10,578 genes and 73 RNAs. *S. thajiwasiensis* ASQP_92 genome was assembled into ∼8.1 Mb having 82 contigs (N50 value of 297,940 and L50 value of 7) with annotation of 7,772 genes and 72 RNAs. For *S. mirabilis* ASQP_98, the assembly was of ∼12 Mb size with 321 contigs (N50 value of 215,380 bp and L50 value of 18) annotated into 11,885 genes and 73 RNAs.

**Table 1:** Genome assembly and annotation features using RAST (Rapid Annotation using Subsystems Technology) analysis.

| Genome and Annotation Statistics |  | *S. violarius*<br>ASQP_29 | *S. rhizosphaerihabitans*<br>ASQP_78 | *S. fulvoviolaceus*<br>ASQP_80 | *S. thajiwasiensis*<br>ASQP_92 | *S. mirabilis*<br>ASQP_98 |
| --- | --- | --- | --- | --- | --- | --- |
| Genome Statistics | Genome size (Total) | 8,909,626 | 10,215,804 | 11,040,768 | 8,133,314 | 12,013,034 |
|  | A | 1,302,791 | 1,510,069 | 1,639,025 | 1,172,073 | 1,797,994 |
|  | T | 1,307,570 | 1,519,768 | 1,643,132 | 1,168,216 | 1,794,744 |
|  | G | 3,153,283 | 3,586,642 | 3,879,410 | 2,896,536 | 4,211,683 |
|  | C | 3,145,982 | 3,599,325 | 3,879,201 | 2,896,489 | 4,208,613 |
|  | N | 0 | 0 | 0 | 0 | 0 |
|  | GC% | 71 | 70 | 70 | 71 | 70 |
|  | AT% | 29 | 30 | 30 | 29 | 30 |
|  | N50 | 27,723 | 147,426 | 69,118 | 297,940 | 215,380 |
|  | L50 | 100 | 23 | 51 | 7 | 18 |
|  | Largest contig size | 114,409 | 424,197 | 291,971 | 1,374,297 | 830,702 |
|  | Smallest contig size | 224 | 201 | 242 | 205 | 203 |
|  | Average contig size | 15,230 | 41,868 | 34,502 | 99,187 | 37,424 |
|  | Total contigs | 585 | 244 | 320 | 82 | 321 |
| Annotation statistics | Total genes | 8,898 | 9,853 | 10,578 | 7,772 | 11,885 |
|  | Coding density | 89% | 85% | 89% | 86% | 86% |
|  | Max gene length | 11,445 | 24,099 | 18,969 | 14,460 | 13,908 |
|  | Min gene length | 60 | 69 | 60 | 75 | 60 |
|  | Average gene length | 898 | 893 | 939 | 915 | 882 |
|  | Max protein length | 3,814 | 8,032 | 6,322 | 4,819 | 4,635 |
|  | Min protein length | 20 | 23 | 20 | 24 | 20 |
|  | Average | 298 | 297 | 312 | 304 | 293 |

| Genome and Annotation Statistics |  | **S.**<br>***violarus***<br>**ASQP_29** | **S.**<br>***rhizosphaerihabita***<br>**ns ASQP_78** | **S.**<br>***fulvoviolace***<br>**us ASQP_80** | **S.**<br>***thajiwasiens***<br>**is ASQP_92** | **S.**<br>***mirabilis***<br>**ASQP_98** |
| --- | --- | --- | --- | --- | --- | --- |
|  | protein length |  |  |  |  |  |
|  | Genes on positive strand | 4,590 | 4,863 | 5,282 | 3,928 | 5,807 |
|  | Genes on negative strand | 4,308 | 4,990 | 5,296 | 3,844 | 6,078 |
|  | Total RNA | 61 | 79 | 73 | 72 | 73 |
|  | Hypothetical proteins | 2,688 | 3,836 | 3,594 | 3,069 | 4,702 |
|  | % of Hypothetical proteins | 30.21 | 38.93 | 33.98 | 39.49 | 39.56 |

All these genomes have 82-585 contigs with different levels of assembly completeness. The genome of *S. thajiwasiensis* ASQP_92 is of the smallest size with the highest assembly quality as we were able to assemble it with an L50 value of 7, the largest contig size of 1.37 Mb, and the average contig size of 99 Kb. The coding density of these strains is also very high ranging from 85-89% as expected. Along with this, another characteristic feature of all five *Streptomyces* strains genomes is the prediction of the exceptionally high number of hypothetical proteins (2,688 in ASQP_29, 3,836 in ASQP_78, 3,594 in ASQP_80, 3,069 in ASQP_92, 4,702 in ASQP_98), which ranges from 30 to 40% of their total coding genes.

### COG (Clusters of Orthologous Groups) analysis classified proteins into different functional categories

The COG (clusters of orthologous groups) classification system was employed to investigate the proteins encoded in the five *Streptomyces* genomes and classify them into functional categories based on their evolutionary relationships. A total number of four main COG categories were classified in each of the five *Streptomyces* species genomes; the highest percentage of COG-categorized genes were assigned to metabolism (39-43%), with a similar proportion distributed across the remaining three categories; 16-18% to cellular processes and signaling, 19-21% to information storage and processing and 20-22% were poorly characterized (Fig.4a). Further, the four main categories were classified into twenty-four COG sub-categories, out of which nine were assigned to cellular processes and signaling, five to information storage and processing, eight to metabolism and two sub-categories were poorly characterized (Fig.4b). A fairly large number of predicted protein-coding genes were classified to secondary metabolite biosynthesis, transport, and catabolism (Q) the highest being in ASQP_98 (389) and ASQP_80 (373). Furthermore, a significant number of genes were assigned to defense mechanisms (V) and posttranslational modification, protein turnover, and chaperones (O) in these antimicrobial-producing *Streptomyces*. Additionally, a significant number of genes had functions that could not be accurately predicted, thus remaining poorly characterized (Fig.4b).

### Gene Ontology highlighted a differential, yet comparable range of GO terms assigned to three different categories

We utilized the standard Gene Ontology vocabulary system to describe three types of Gene Ontology terms: molecular function, biological pathway or larger process, and cellular location for genes across all five *Streptomyces* genomes. The percentage of total RAST annotated genes assigned to different GO terms ranged from 21-26% across different *Streptomyces* strains (Fig.5a). The maximum percentage of GO terms were assigned to molecular function (49-50%), followed by biological processes (36-37%) and cellular component (13-14%) (Fig.5b). Although, in each *Streptomyces* strain the GO terms were assigned differently to three different categories, the assignment of a particular category to all the five strains exhibited some degree of similarity.

### Pfam analysis revealed that the average top Pfam domains are involved in several antimicrobial-related biological processes

In this study, we conducted an analysis to determine the number and uniqueness of protein family (Pfam) domains among all the five *Streptomyces* species. We then assessed the prevalence of these unique domains in each species and calculated the mean value across all observed protein family domains. The total number of unique Pfam domains in each strain was 9,047 in *S. violarus* ASQP_29, 9,354 in *S. rhizosphaerihabitans* ASQP_78, 10,922 in *S. fulvoviolaceus* ASQP_80, 7,626 in *S. thajiwasiensis* ASQP_92 and 11,361 in *S. mirabilis* ASQP_98 (Fig.5c). A total of 6238 proteins were identified to have Pfam domains in ASQP_29, 6377in ASQP_78, 7522 in ASQP_80, 5226 in ASQP_92, and the highest count being 7770 in ASQP_98.

The top 30 domains were then identified, and their average abundance was reported (Fig.5d). Among these top 30 domains, we could identify protein family domains involved in several biological processes, including 1) domains that function as part of transport proteins, 2) DNA binding domains, 3) domains as part of several enzymes like dehydrogenases, transferases, and hydrolases and 4) domains involvedin signal transduction.

#### Transport Pfam domains

Among the top 30 Pfam domains, we identified several transport proteins including ABC_trans (ATP binding cassette transporter), BPD_transp_1 (binding protein-dependent transporter system1) MFS_1 (major facilitator superfamily transporter), HATPases (histidine kinase-like ATPases) like HATPase_C, HATPase_c_2, which are important components of ABC transporters. These transporters are involved in the uptake and efflux of a wide range of compounds including nutrients, amino acids, peptides, signaling molecules, secondary metabolites, and antibiotics. The intake, efflux, and translocation of these substrates and metabolites across biological membranes facilitate the growth, development, and defense and are important for antibiotic biosynthesis, morphological differentiation, and antimicrobial resistance (Crits-Christoph et al., 2021; Méndez & Salas, 2001; Yang et al., 2022; Zhou et al., 2016).

#### DNA binding domains

The protein family domains that are included in this category are TetR_N (Tetracycline repressor N terminal domain) GerE (bacterial spore germination domain), Helix turn Helix domain like HTH_31 and HTH_1, GntR (Gluconate operon repressor, Trans_reg_C (C terminal DNA binding domain of bacterial transcriptional regulators), LysR_substrate (Lysine specific regulatory proteins), Sigma factors domains of bacterial RNA polymerase like sigma70_r2 and sigma70_r4_2. These DNA binding domains act as transcription regulators and perform multiple functions that include antibiotic resistance (TetR_N), late-stage development and sporulation in *Streptomyces* (GerE), aerial mycelial formation and sporulation (HTH_1 and HTH_31), regulation of secondary metabolism (GntR, Trans_reg_C), quorum sensing, virulence and stress response (LysR) and domains involved in transcriptional initiation and proper response to oxidative stress (sigma70_r2 and sigma70_r4_2). Thus, most of the domains in this category are involved in growth, morphological differentiation and sporulation, antibiotic resistance, and regulation of oxidative stress. All these processes are directly or indirectly involved in secondary metabolite regulation and antibiotic biosynthesis.

#### Domains of signal transduction pathways

These include response_reg (response regulator)- a part of bacterial two-component regulation system, Kinases such as HisKA_3 (histidine kinase domain) and Pkinase (protein kinase). These domains play a crucial role in the activation and inhibition of downstream response regulators and regulation of different processes including secondary metabolism and antibiotic biosynthesis (Cruz-Bautista et al., 2023; Mikulík et al., 2002; Ni et al., 2019; Otten et al., 1995). We also performed specific domain analysis to find the chemosensory systems and as expected, we were not able to identify any chemosensory components in these organisms (Sharma, Khatri, et al., 2018; Sharma, Parales, et al., 2018; Zhu et al., 2023).

#### Enzyme domains

The domains included in this category form a structural part of several enzymes like dehydrogenases, transferases, and hydrolases. The dehydrogenases include domains of alcohol dehydrogenases (ADHs) like Adh_shortand Adh_short_C2 (short-chain alcohol dehydrogenase and C terminal domain of short-chain alcohol dehydrogenase respectively), ADH_N (conserved domain of 70-80 amino acids located at the N terminal of various alcohol dehydrogenases), ADH_Z_N (Zinc containing domain binding to N terminal of alcohol dehydrogenase) and acyl-CoA dehydrogenase domains (Acyl-CoA_dh_N and Acyl-CoA_dh_1).The transferase and hydrolase identified include Acetyltransf_1(acetyl transferase_1) and Abhydrolase_1 (a domain that belongs to the α/β-hydrolase family of enzymes). These enzymes are involved in reactions like oxidation and reduction of alcohols, aldehydes, and other substrates (e.g., ADH-alcohol dehydrogenases), acetyl group transfer from acetyl-CoA to a variety of substrates including amino acids, peptides, and small molecules (Acetyltransf_1) and hydrolysis, esterification and transesterification reactions (Abhydrolase_1). Most of these enzymes and the reactions they catalyze are central to the secondary metabolite biosynthesis and regulation by PKS and NRPS and these enzymes play an important role in antibiotic and bioactive metabolite biosynthesis (Aparicio et al., 1996; Jenke-Kodama & Dittmann, 2009)

In addition to these Pfam domains, we were able to identify adenosine mono phosphate binding (AMP-binding and AMP-binding_C) and SPOIIE (stage O sporulation protein IIE) domains. AMP binding domain binds and senses the intracellular concentration of AMP and thus modulates the intracellular activities of several enzymes in response to these AMP levels, thereby controlling several processes including antibiotic production (Latoscha et al., 2019). SPOIIE domain plays a role in switching from vegetative growth to sporulation in response to adverse environmental conditions (Miguélez et al., 1998; Schwedock et al., 1997; Sharp & Pogliano, 2003).

### Gene mapping of protein-expressed genes (PEGs) predicted their varied assignment to different functional categories

We mapped the percentage of predicted protein-expressed genes (PEGs) to different functional categories across the five *Streptomyces* genomes (Fig. 5e). The percentage of predicted protein-coding genes that were classified into different COG functional categories ranged from 72-80%; 78% in *S. violarus* ASQP_29, 71% in *S. rhizosphaerihabitans* ASQP_78, 80% in *S. fulvoviolaceus* ASQP_80, 74% in *S. thajiwasiensis* ASQP_92 and 72% in *S. mirabilis* ASQP_98. Further, the percentage of total genes assigned to gene ontology terms ranged from 22-26% and the value of gene ontologies in these *Streptomyces* were 26% in *S. violarus* ASQP_29 and *S. mirabilis* ASQP_98, 24% in *S. rhizosphaerihabitans* ASQP_78, 27% in *S. fulvoviolaceus* ASQP_80 and 22% in *S. thajiwasiensis* ASQP_92.

**Figure 5.**
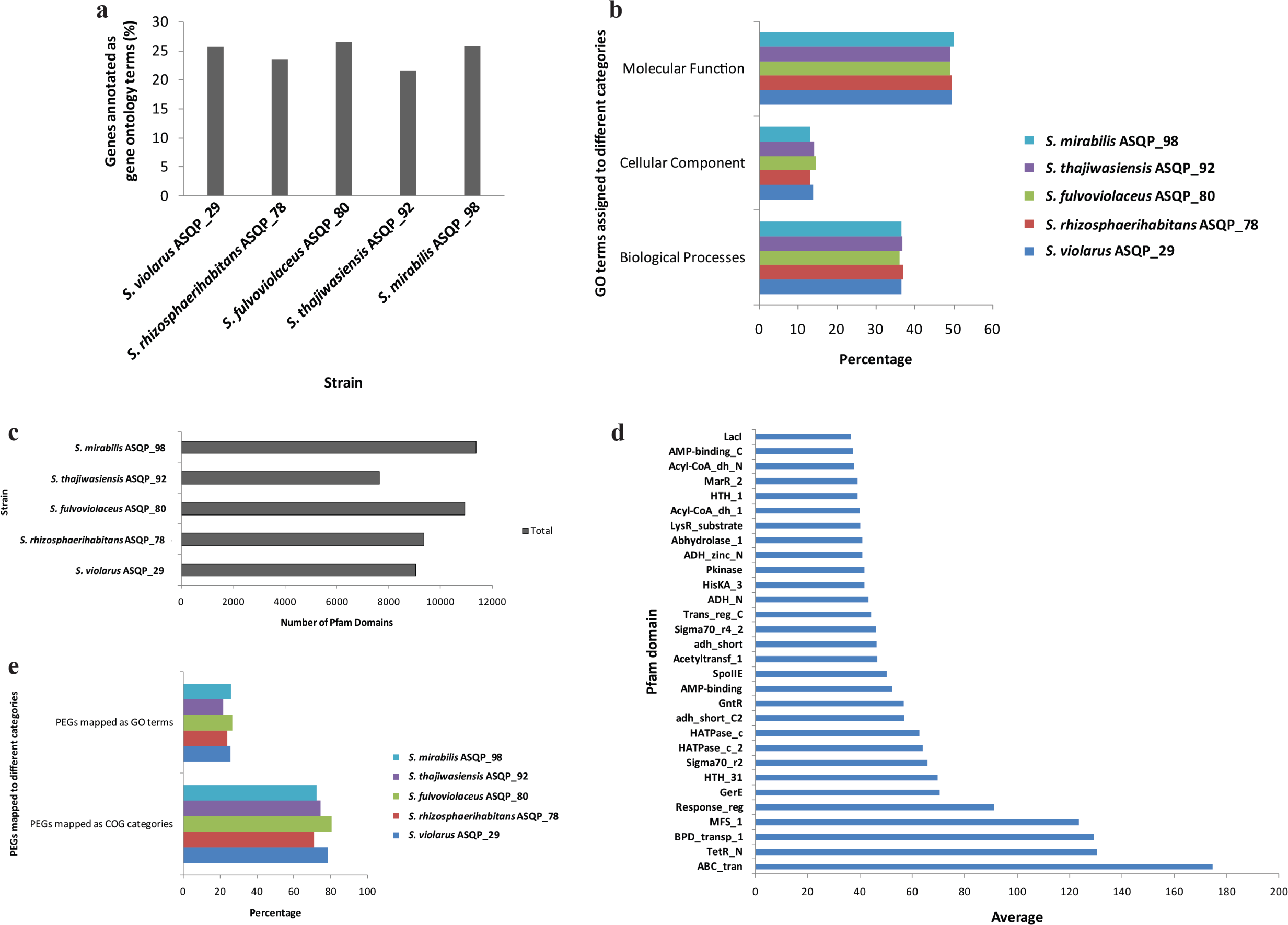
Gene ontology (GO) annotation, protein family domain (Pfam) analysis, and protein expression genes (PEGs) mapping among five *Streptomyces* species. a) Percentage of total genes assigned to gene ontology terms. b) Percentage of GO terms assigned to three different categories. c) Number of unique Pfam domains in each *Streptomyces* species. d) Average of the top 30 Pfam domains among five *Streptomyces* species. e) Percentage of PEGs mapped to different COG categories and GO terms.

### Biosynthetic Gene Clusters (BGCs) classification and profiling of assembled bioactive *Streptomyces* displayed the encoding of conserved BGCs, strain-specific BGCs, low similarity percentage BGCs, and orphan BGCs

We analyzed the genomes of five bioactive *Streptomyces* against antiSMASH using the MiBiG database. This allowed us to identify both conserved and strain-specific BGCs, which we then classified into different categories. A total of 182 BGCs were classified in all the five *Streptomyces* species; 47 in *Streptomyces violarus* ASQP_29, 34 each in *Streptomyces rhizosphaerihabitans* ASQP_78 and *Streptomyces fulvoviolaceus* ASQP_80, 32 in *Streptomyces thajiwasiensis* ASQP_92 and 35 in *Streptomyces mirabilis* ASQP_98 (Fig. 6a).Thus, irrespective of their low genome sizes, the maximum number of BGCs was reported in *Streptomyces violarus* ASQP_29 and *Streptomyces thajiwasiensis* ASQP_92. The BGCs that are abundantly present and conserved in all five *Streptomyces* include Polyketide Synthases (PKS), Non-Ribosomal Peptide Synthases (NRPS), terpenes, siderophores, and RiPP-like BGCs, however other BGCs that are also represented in all the five *Streptomyces* albeit less abundantly include NAPAA, melanin, lanthipeptides, and ectoines (Fig. 6b). Further, a substantial number of BGCs were reported as Hybrid categories; these include NRPS-PKS hybrid, NRPS hybrids with NAPAA, oligosaccharides, phosphonates and RREcontaining BGCs; PKS hybrids with oligosaccharides, indole, hgIE and thioamides and terpene hybrids with butyrolactones, lanthipeptides, and melanins. Also, some NRPS and PKS form hybrids with categories not assigned to any BGC class (hybrids with class other) (Fig. 6b). Despite the conserved BGCs present across five *Streptomyces* strains like NRPS, PKS, ectoines, lanthipeptides, melanin, siderophores etc., we could observe one or multiple strain specific BGCs in four of the five strains; these strain specific BGCs include one hybrid BGC in *Streptomyces violarus* ASQP_29 (T3PKS,thioamitides), two hybrids in *Streptomyces fulvoviolaceus* ASQP_80 (terpene,butyrolactone and terpene,melanin), seven in *Streptomyces thajiwasiensis* ASQP_92 out of which five are hybrid BGCs (betalactone,hglE-KS; NRPS,oligosaccharide,T2PKS,PKS-like,RRE-containing; NRPS,RRE-containing,phosphonate,T1PKS,NRPS-like; other,terpene and thiopeptide,RiPP-like,thioamide-NRP,ladderane,NRPS) and two are non-hybrid BGCs (phenazine and RRE-containing) and finally four in *Streptomyces mirabilis* ASQP_98 out of which three are hybrid BGCs (NRPS,NAPAA; T2PKS,indole and terpene,lanthipeptide-class-ii) and one is non hybrid (lassopeptide) (Fig. 6a and 6b). The minimum number of total BGCs (32) and the maximum number of strain-specific BGCs (7) in *Streptomyces thajiwasiensis* ASQP_92 further stressed the novel taxonomic and biosynthetic character of this strain. Although the presence of conserved BGCs across these *Streptomyces* strains specifies their evolutionary relatedness in the production of different antimicrobial secondary metabolites, the strain-specific encoding of BGCs highlights their significance in fine-tuning the metabolic potential of a particular strain in response to specific environmental stimuli and may represent as novel metabolites within these antimicrobial secretion bioactive strains (Otani et al., 2022).

**Figure 6.**
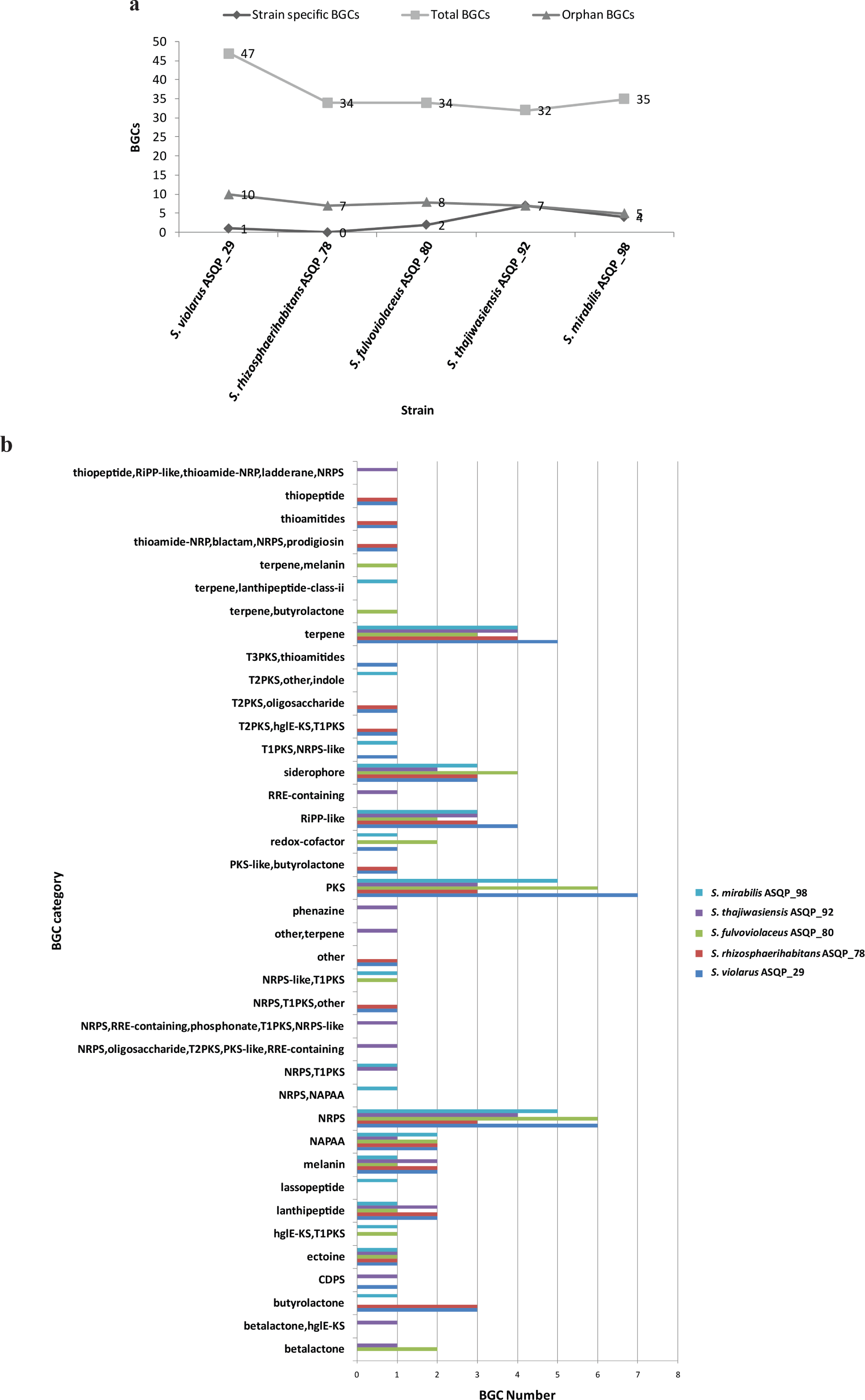
BGC (Biosynthetic Gene Cluster) analysis of five *Streptomyces* species. a) Number of total, strain-specific and orphan BGCs. b) The classification of BGCs, their profiling into different categories, and the value of each BGC category against five *Streptomyces* species.

Looking into the similarity characteristics of BGCs to the known BGC clusters, we could identify several BGCs whose percentage similarity ranged between 60-100% to those of known secondary metabolites. These highly similar metabolites among others include ectones, melanin, lipopeptides, mycobactin desferrioxamies, saprolmycin E, germicidin, resorsinols, collinomycin, gamma butyrolactones,andradamycin. The full spectrum of BGCs and their percentage similarity is shown in Supplementary Table 4. Out of a total of 182 BGCs in these five bioactive *Streptomyces*, 27 different clusters (out of which ten in *Streptomyces violarus* ASQP_29, seven each in *S. rhizosphaerihabitans* ASQP_78 and *S. thajiwasiensis* ASQP_92, eight in *S. fulvoviolaceus* ASQP_80, five in *S. mirabilis* ASQP_98) could be identified with no known homologs and are classified as orphan BGCs (Fig. 6a and Supplementary table 4). The presence of orphan BGCs and BGCs with low similarity values to a particular known metabolite cluster further signifies the novel encoding biosynthetic potential in these *Streptomyces* strains (Malik et al., 2020; Shi et al., 2019).

## Discussion

The use of both 16S rDNA and gyrB phylogenetic analyses for five bioactive *Streptomyces* species provided valuable information about their evolutionary history and allowed for a more comprehensive understanding of the genetic relatedness among these species. The results supported the usage of different alternative molecular markers for the accurate identification of *Streptomyces* species (Case et al., 2007; Witt & Stackebrandt, 1990). Furthermore, the study provides valuable insights into the genome-based taxonomic classification (phylogenomic approach) and highlights the usefulness of combining multiple methods for robust classification as performed earlier too (Mohammadipanah & Dehhaghi, 2017; van der Aart et al., 2019).

One notable feature of the genome annotation statistics of all five *Streptomyces* strains is the prediction of a high number of hypothetical proteins, with ASQP_78 exhibiting the highest number of 3,836 hypothetical proteins. These proteins lack known functional domains or sequence similarity to known proteins, making their functions difficult to predict. The presence of a large number of hypothetical proteins suggests that there is still much to be discovered about the genomes of these *Streptomyces* strains and their potential for biotechnological applications (Ferdous et al., 2020; Tierrafría et al., 2016). Similarly, our COG and GO analysis revealed significant numbers of genes whose function could not be predicted properly and thus remained poorly characterized. It is important to note that poorly characterized genes may hold significant importance in the adaptation and survival of these *Streptomyces* species in their natural habitats, and further research is necessary to understand their functions (Galperin et al., 2015; Tatusov et al., 1997). The significant number of genes associated with secondary metabolite biosynthesis and defense mechanisms in these *Streptomyces* species suggests their potential as sources of natural products with antimicrobial properties. Furthermore, the differences in the number of genes assigned to COG categories among the five strains may be due to differences in their genome sizes.

The Pfam domains study provides a comprehensive analysis of the protein family domains in *Streptomyces* species and their roles in various biological processes, including secondary metabolite and antibiotic biosynthesis, antibiotic resistance, nutrient uptake and efflux, late-stage development and sporulation, defense and regulation of oxidative stress. These findings could be valuable for understanding the genetic basis of the diverse physiological and ecological roles of *Streptomyces* and for developing new strategies for the discovery of bioactive compounds (Romero-Rodríguez et al., 2015; Zhou et al., 2016).

BGCs that show a low similarity percentage to known metabolites in antiSMASH are often encoded with a repertoire of cryptic regulatory genetic elements as their products have not yet been characterized or identified. The identification and activation of these cryptic regulatory elements in BGCs could enhance the production of their encoded metabolites. The researchers have thus focused on identifying and manipulating regulatory genes that encode proteins that act as transcription factors, DNA-binding proteins, and other regulatory proteins that control the expression of genes within BGCs (Bc et al., 2021; Rutledge & Challis, 2015). The presence of different strain-specific BGCs highlights the significance of the selective nature of different *Streptomyces* in the production of different bioactive metabolites (Otani et al., 2022). Further, the discovery and characterization of orphan BGCs in *Streptomyces* have significant implications for drug discovery and development. Our lab is constantly putting efforts into synthesizing these BGCs as they can potentially produce new compounds with antimicrobial, anticancer, or other pharmacological activities, and may serve as new leads for drug development. Additionally, more research on these orphan BGCs and their regulation can provide insight into the evolution of natural product biosynthesis and gene regulation in *Streptomyces* (Amos et al., 2017; Rutledge & Challis, 2015; van der Heul et al., 2018).

## Conclusion

This comparative genomics study of five bioactive *Streptomyces* species isolated from NWH has significantly imparted to our knowledge of their taxonomic classification and stressed the use of multiple molecular markers and polyphasic approach for genetic relatedness and accurate species identification. The genome annotation and functional categorization revealed the identification of a substantial number of hypothetical proteins and the presence of poorly characterized genes which underscores the complexity of *Streptomyces* genomes and proposes the need for further research for bioactive compound elucidation and biotechnological exploration of these microorganisms. Furthermore, the study highlighted the diverse repertoire of BGCs, including conserved, strain-specific, and orphan BGCs with no known homologs, and stressed the potential of *Streptomyces* as rich biosynthetic producers and sources of novel bioactive compounds. Overall, this study contributes to our understanding of *Streptomyces* species elucidation, genomic complexity, functional annotation, and biosynthetic capabilities, thus paving the way for future research in antibiotic discovery and natural product biosynthesis. The extensive genomic research conducted on the *Streptomyces* of high altitudes of NWH will open the possibilities of further diversity exploration, species identification, functional prediction, biosynthetic exploitation, and novel bioactive metabolite characterization within the different species of this genus.

## Conflict of Interest Statement

The authors declare that the work reported in this paper is devoid of any known conflicting financial interests or personal relationships.

## Acknowledgments

The authors in this manuscript would like to thank CSIR-Indian Institute of Integrative Medicine (IIIM), Jammu; NCMR-National Centre for Cell Science (NCCS), Pune for providing facilities and space in the laboratory. We acknowledge the University Grants Commission (UGC) for providing a fellowship to Aasif Majeed during the tenure of this work. Mariam Azeezuddin Haneen is supported by the DST-INSPIRE Research fellowship. Gaurav Sharma acknowledges the DST-INSPIRE Faculty Award from the Government of India and a seed grant from the Indian Institute of Technology, Hyderabad.

## Availability of data and materials

The whole-genome sequence (WGS) assembled files used in this study are available at NCBI’s Genome portal with accession numbers PRJNA879315 for *S. violarus* ASQP_29, PRJNA879313 for *S. rhizosphaerihabitans* ASQP_78, PRJNA879311 for *S. fulvoviolaceus* ASQP_80, PRJNA879322 for *S. thajiwasiensis* ASQP_92 and PRJNA879317 for *S. mirabilis* ASQP_98.

The pure cultures of these strains are deposited in the Microbial Culture Collection (MCC) repository at National Centre for Microbial Resources (NCMR), Pune, India under accession numbers MCC 5213 for *S. violarus* ASQP_29, MCC 5214 for *S. rhizosphaerihabitans* ASQP_78, MCC 5215 for *S. fulvoviolaceus* ASQP_80, MCC 5216 for *S. thajiwasiensis* ASQP_92, and MCC 5217 for *S. mirabilis* ASQP_98. NCBI will update the name of *S. thajiwasiensis* sp. nov. ASQP_92 after the peer review and acceptance of this manuscript; till then they have kept the name as *Streptomyces* sp. ASQP_92.

## Supplementary Data Legends

**Supplementary Table 1:** Average nucleotide identity based on MUMmer (ANIm) genome characteristics of query *Streptomyces* strains and their closely related type strains

**Supplementary Table 2:** Digital DNA-DNA hybridization using Genome to Genome Distance Calculator (GGDC)

**Supplementary Table 3**: Pairwise digital DNA-DNA Hybridization (dDDH) values between the query genomes and the top type stains genomes using Type Strain Genome Server (TYGS).

* Three different Genome BLAST Distance Phylogeny (GBDP) formulas along with the confidence intervals (C.I.) for the most recommended formula (*d_4_)* used for phylogenomic analyses are:

- formula *d_0_* (a.k.a. GGDC formula 1): length of all HSPs (high-scoring segment pair) divided by total genome length
- formula *d_4_* (a.k.a. GGDC formula 2): sum of all identities found in HSPs divided by overall HSP length (**most recommended formula for incomplete draft genomes**)
- formula *d_6_* (a.k.a. GGDC formula 3): sum of all identities found in HSPs divided by total genome length

**Supplementary Table 4:** BGC type with percentage similarity to the most similar known cluster.

**-** absent, **+** present, values in parenthesis are percentage values to the most similar known cluster. Number of orphan type BGC clusters when present in more than one in a particular *Streptomyces* species are also specified after the + sign.

## Notes

### Competing Interest Statement

The authors have declared no competing interest.

