## Supplementary Table 1 for "Comparative genomics studies provide insights into the taxonomic classification and secondary metabolic potential of five bioactive *Streptomyces* species isolated from the North-Western Himalaya"

**Supplementary Table 1:**Average nucleotide identity based on MUMmer (ANIm) genome characteristics of query Streptomyces strains and their closely related type strains.

| Query genome | Reference genome | ANIm (%) |
|---|---|---|
| ASQP_29 | *Streptomyces violarus* JCM4534 | 0.831 |
| | *Streptomyces janthinus* JCM4387 | 0.807 |
| | *Streptomyces purpurascens* JCM4509 | 0.767 |
| ASQP_78 | *Streptomyces populi* A249 | 0.660 |
| | *Streptomyces triticiradicis* NEAU-H2 | 0.651 |
| | *Streptomyces aureus* NRRLB-1941 | 0.628 |
| ASQP_80 | *Streptomyces fulvoviolaceus* NRRLB-2870 | 0.880 |
| | *Streptomyces lincolnensis* LC-G | 0.605 |
| | *Streptomyces resistomycificus* DSM40133 | 0.599 |
| | *Streptomyces pseudovenezuelae* DSM40212 | 0.589 |
| ASQP_92 | *Streptomyces hundungensis* BH38 | 0.785 |
| | *Streptomyces xanthochromogenes* JCM4594 | 0.617 |
| | *Streptomyces violascens* NBRC12920 | 0.588 |
| ASQP_98 | *Streptomyces mirabilis* JCM4551 | 0.805 |
| | *Streptomyces olivochromogenes* NBRC3561 | 0.787 |
| | *Streptomyces fagopyri* QMT-28 | 0.617 |
