## Supplementary Table 2 for "Comparative genomics studies provide insights into the taxonomic classification and secondary metabolic potential of five bioactive *Streptomyces* species isolated from the North-Western Himalaya"

**Supplementary Table 2**: Digital DNA DNA hybridization using Genome to Genome Distance Calculator (GGDC)

| Query genome | Reference genome | % dDDH (Formula 2*) | Model C.I. | Distance | Prob. DDH > = 70 % | G+C difference |
|---|---|---|---|---|---|---|
| **ASQP_29** | *Streptomyces janthinus* ASM1464967v1 | 77.1 | [74.1 - 79.8%] | 0.0268 | 88.04 | 0.09 |
| | *Streptomyces violarus* ASM1465025v1 | 71.8 | [68.8 - 74.6%] | 0.0337 | 81.33 | 0.18 |
| | *Streptomyces purpurascens* ASM1465015v1 | 62.7 | [59.8 - 65.5%] | 0.0471 | 60.88 | 0.31 |
| **ASQP_78** | *Streptomyces populi* ASM291101v1 | 40.4 | [37.9 - 43%] | 0.098 | 3.02 | 1.32 |
| | *Streptomyces triticiradicis* ASM886868v1 | 40.4 | [37.9 - 42.9%] | 0.0981 | 2.99 | 1.15 |
| | *Streptomyces olivochromogenes* ASM151411v1 | 30.4 | [28 - 32.9%] | 0.1401 | 0.12 | 0.11 |
| | *Streptomyces avermitis* ASM151413v1 | 28.3 | [25.9 - 30.8%] | 0.1521 | 0.05 | 0.4 |
| | *Streptomyces aureus* ASM141813v1 | 23.5 | [21.2 - 25.9%] | 0.1862 | 0 | 1.51 |
| **ASQP_80** | *Streptomyces fulvoviolaceus* ASM71816v1 | 94.8 | [93.2 - 96.1%] | 0.0069 | 97.15 | 0.02 |
| | *Streptomyces resistomycificus* ASM71621v1 | 31.3 | [28.9 - 33.8%] | 0.1354 | 0.18 | 0.6 |
| | *Streptomyces canus* ASM151414v1 | 31.3 | [28.9 - 33.8%] | 0.1354 | 0.18 | 0.03 |
| | *Streptomyces canus* ASM151404v1 | 31.3 | [28.9 - 33.9%] | 0.135 | 0.180 | 0.070 |
| | *Streptomyces resistomycificus* ASM151426v1 | 30.9 | [28.5 - 33.4%] | 0.1373 | 0.15 | 0.77 |
| | *Streptomyces resistomycificus* ASM127050v1 | 30.9 | [28.5 - 33.4%] | 0.1371 | 0.15 | 0.76 |
| | *Streptomyces pseudovenezuelae* ASM151395v1 | 30.7 | [28.3 - 33.2%] | 0.138 | 0.140 | 0.750 |
| | *Streptomyces lincolnensis* ASM168535v1 | 30.4 | [28 - 32.9%] | 0.1398 | 0.13 | 0.74 |

| **Query genome** | **Reference genome** | **% dDDH (Formula 2*)** | **Model C.I.** | **Distance** | **Prob. DDH > = 70 %** | **G+C difference** |
|---|---|---|---|---|---|---|
| | *Streptomyces chartreusis* ASM870471v1 | 27.9 | [25.6 - 30.4%] | 0.1541 | 0.04 | 0.72 |
| | *Streptomyces triticiradicis* ASM886868v1 | 25.9 | [23.6 - 28.4%] | 0.1676 | 0.01 | 1.22 |
| **ASQP_92** | *Streptomyces hundungensis* ASM362781v1 | 49.9 | [47.2 - 52.5%] | 0.0719 | 18.74 | 0.29 |
| | *Streptomyces violascens* ASM1464995v1 | 36.3 | [33.9 - 38.8%] | 0.1128 | 0.99 | 0.75 |
| | *Streptomyces xanthochromogenes* ASM1465035v1 | 36.3 | [33.9 - 38.8%] | 0.1128 | 0.99 | 0.04 |
| | *Streptomyces xanthochromogenes* ASM1465045v1 | 36.2 | [33.8 - 38.7%] | 0.1133 | 0.95 | 0.12 |
| | *Streptomyces melanogenes* ASM1464979v1 | 25.7 | [23.4 - 28.2%] | 0.169 | 0.01 | 0.12 |
| | *Streptomyces chryseus* ASM1465075v1 | 23.2 | [20.9 - 25.7%] | 0.1885 | 0 | 0.37 |
| **ASQP_98** | *Streptomyces mirabilis* ASM1465027v1 | 68.1 | [65.1 - 70.9%] | 0.0388 | 74.57 | 0.18 |
| | *Streptomyces olivochromogenes* ASM151411v1 | 68.1 | [65.1 - 70.9%] | 0.0389 | 74.51 | 0.13 |
| | *Streptomyces fagopyri* ASM949827v1 | 33.0 | [30.5 - 35.5%] | 0.1272 | 0.33 | 1.15 |
| | *Streptomyces avermitilis* (high GC Gram+) ASM540598v1 | 29.7 | [27.3 - 32.2%] | 0.1441 | 0.09 | 0.44 |

*Recommended formula (identities sum in high-scoring segment pairs (HSPs) or maximally unique matches (MUMs)/ overall HSP length) used against incomplete draft genomes.
