## Supplementary Table 3 for "Comparative genomics studies provide insights into the taxonomic classification and secondary metabolic potential of five bioactive *Streptomyces* species isolated from the North-Western Himalaya"

**Supplementary Table 3**: Pairwise digital DNA-DNA Hybridization (dDDH) values between the query genomes and the top type stains genomes using Type Strain Genome Server (TYGS).

* Three different Genome BLAST Distance Phylogeny (GBDP) formulas along with the confidence intervals (C.I.) for the most recommended formula (*$d_4$*) used for phylogenomic analyses are:

- formula $d_0$ (a.k.a. GGDC formula 1): length of all HSPs (high-scoring segment pair) divided by total genome length
- formula $d_4$ (a.k.a. GGDC formula 2): sum of all identities found in HSPs divided by overall HSP length (**most recommended formula for incomplete draft genomes**)
- formula $d_6$ (a.k.a. GGDC formula 3): sum of all identities found in HSPs divided by total genome length

| **Query genome** | **Reference genome** | **% dDDH (Formula $d_4$*)** | **Model C.I. ($d_4$)** | **% dDDH (Formula $d_0$*)** | **% dDDH (Formula $d_6$*)** | **G+C difference** |
|---|---|---|---|---|---|---|
| **ASQP_29** | *Streptomyces janthinus* JCM 4387 | 77.1 | [74.1 - 79.8%] | 73.4 | 76.7 | 0.09 |
| | *Streptomyces violarus* CECT 3237 | 71.8 | [68.8 - 74.6%] | 76.0 | 77.9 | 0.19 |
| | *Streptomyces violarus* JCM 4534 | 71.8 | [68.8 - 74.6%] | 76.0 | 77.9 | 0.18 |
| | *Streptomyces purpurascens* JCM 4509 | 62.7 | [59.8 - 65.5%] | 66.3 | 67.5 | 0.39 |
| **ASQP_78** | *Streptomyces triticiradicis* NEAU-H2 | 40.4 | [37.9 – 42.9%] | 46.7 | 45.0 | 1.15 |
| | *Streptomyces populi* A249 | 40.4 | [37.9 - 43.0%] | 46.4 | 44.8 | 1.32 |
| | *Streptomyces albiflavescens* CGMCC4.7111 | 32.1 | [29.7 - 34.6%] | 33.0 | 31.7 | 0.39 |
| | *Streptomyces dengpaensis* XZHG99 | 31.5 | [29.1 - 34.0%] | 30.0 | 29.0 | 0.42 |
| | *Streptomyces olivochromogenes* DSM40451 | 30.3 | [27.9 - 32.8%] | 30.4 | 29.2 | 0.11 |
| | *Streptomyces avermitilis ATCC 31267* | 28.3 | [26.0 - 30.8%] | 28.8 | 27.5 | 0.19 |

| Query genome | Reference genome | % dDDH (Formula $d_4$*) | Model C.I. ($d_4$) | % dDDH (Formula $d_0$*) | % dDDH (Formula $d_6$*) | G+C difference |
|---|---|---|---|---|---|---|
| ASQP_80 | *Streptomyces resistomycificus* NRRL ISP-5133 | 31.3 | [28.9 - 33.8%] | 34.7 | 32.9 | 0.60 |
| | *Streptomyces canus* DSM 40017 | 31.3 | [28.9 - 33.8%] | 37.8 | 35.4 | 0.02 |
| | *Streptomyces ciscaucasicus* DSM40275 | 31.3 | [28.9 - 33.8%] | 38.9 | 36.3 | 0.08 |
| | *Streptomyces resistomycificus* NRRL 2290 | 30.9 | [28.5 - 33.4%] | 37.7 | 35.2 | 0.77 |
| | *Streptomyces resistomycificus* DSM 40133 | 30.9 | [28.5- 33.4%] | 37.7 | 35.2 | 0.78 |
| | *Streptomyces pseudovenezuelae* DSM 40212 | 30.7 | [28.3 - 33.2%] | 38.8 | 36.1 | 0.76 |
| | *Streptomyces lincolnensis* NRRL 2936 | 30.4 | [28.0 - 32.9%] | 39.4 | 36.4 | 0.74 |
| | *Streptomyces phaeopurpureus* DSM 40125 | 30.0 | [27.6 - 32.5%] | 36.1 | 33.7 | 0.71 |
| | *Streptomyces chartreusis* ATCC 14922 | 27.9 | [25.6 - 30.4%] | 34.3 | 31.8 | 0.72 |
| ASQP_92 | *Streptomyces violascens* JCM 4424 | 36.3 | [33.9 - 38.8%] | 42.6 | 40.5 | 0.75 |
| | *Streptomyces michiganensis* JCM4594 | 36.3 | [33.9 - 38.8%] | 46.6 | 43.9 | 0.04 |
| | *Streptomyces xanthochromogenes* JCM4612 | 36.2 | [33.8 - 38.7%] | 46.5 | 43.7 | 0.11 |
| | *Streptomyces melanogenes* JCM 4398 | 25.7 | [23.4 - 28.2%] | 28.2 | 26.6 | 0.13 |
| | *Streptomyces genisteinicus* CRPJ-33 | 23.5 | [21.2 - 26.0%] | 21.0 | 20.4 | 2.21 |
| | *Streptomyces lateritius* JCM 4389 | 23.5 | [21.2 - 25.9%] | 22.8 | 21.8 | 0.12 |
| | *Streptomyces chryseus* JCM 4737 | 23.2 | [20.9 - 25.7%] | 23.9 | 22.7 | 0.37 |
| ASQP_98 | *Streptomyces mirabilis* JCM 4551 | 68.1 | [65.1 - 70.9%] | 65.7 | 68.1 | 0.18 |

| **Query genome** | **Reference genome** | **% dDDH (Formula $d_4$*)** | **Model C.I. ($d_4$)** | **% dDDH (Formula $d_0$*)** | **% dDDH (Formula $d_6$*)** | **G+C difference** |
|---|---|---|---|---|---|---|
| | *Streptomyces olivochromogenes* DSM40451 | 68.0 | [65.1 - 70.9%] | 64.3 | 66.8 | 0.14 |
| | *Streptomyces fagopyri* QMT-28 | 33.0 | [30.5 - 35.5%] | 35.6 | 34.0 | 1.15 |
| | *Streptomyces albiflavescens* CGMCC 4.7111 | 32.0 | [29.6 - 34.5%] | 33.6 | 32.1 | 1.15 |
| | *Streptomyces dengpaensis* XZHG99 | 31.4 | [29.0 - 33.9%] | 29.0 | 28.2 | 0.17 |
| | *Streptomyces avermitilis* (high GC Gram+) ASM540598v1 | 29.7 | [27.3 - 32.2%] | 32.3 | 30.6 | 0.44 |
