## Supplementary Table 4 for "Comparative genomics studies provide insights into the taxonomic classification and secondary metabolic potential of five bioactive *Streptomyces* species isolated from the North-Western Himalaya"

**Supplementary Table 4:** BGC type with percentage similarity to most similar known cluster.

- absent , + present, values in parenthesis are percentage values to most similar known cluster. Number of orphan type BGC clusters when present in more than one in a particular *Streptomyces* species are also specified after the + sign.

| BGC Type | Most similar known cluster type | ASQP_29 | ASQP_78 | ASQP_80 | ASQP_92 | ASQP_98 |
|---|---|---|---|---|---|---|
| **betalactone** | Orphan | - | - | + | + | - |
| **betalactone** | thioviridamide | - | - | + (10) | - | - |
| **betalactone,hglE-KS** | aurantimycin A | - | - | - | + (6) | - |
| **butyrolactone** | Orphan | + | + | - | - | - |
| **butyrolactone** | cyphomycin | + (5) | + (5) | - | - | - |
| **butyrolactone** | griseoviridin / fijimycin A | + 5 | + (5) | - | - | - |
| **butyrolactone** | prejadomycin / rabelomycin / gaudimycin C / gaudimycin D / UWM6 / gaudimycin A | - | - | - | - | + (100) |
| **CDPS** | Orphan | + | - | - | - | - |
| **CDPS** | BD-12 | - | - | - | + (14) | - |
| **ectoine** | ectoine | + (100) | + (100) | + (100) | + (100) | + (100) |
| **hglE-KS,T1PKS** | laidlomycin | - | - | + (6) | - | - |
| **hglE-KS,T1PKS** | toxoflavin / fervenulin | - | - | - | - | + (10) |
| **lanthipeptide-class-i** | Griselimycin | + (7) | + (7) | - | - | - |
| **lanthipeptide-class-ii** | Orphan | - | - | - | + | - |
| **lanthipeptide-class-iii** | Informatipeptin | + (71) | + (71) | + (42) | - | - |
| **lanthipeptide-class-iii** | AmfS | - | - | - | - | + (80) |
| **lanthipeptide-class-iv** | Venezuelin | - | - | - | +(100) | - |
| **lassopeptide** | siamycin I | - | - | - | - | + 56 |
| **melanin** | Istamycin | + (5) | + (5) | + (5) | + (7) | - |
| **melanin** | melanin | + (71) | + (71) | - | + (28) | + (57) |
| **NAPAA** | belactosin A / belactosin C | + (8) | + (8) | - | - | - |
| **NAPAA** | stenothricin | + (13) | + (13) | + (13) | - | + (13) |
| **NAPAA** | Orphan | - | - | + | - | + |
| **NAPAA** | rifamorpholine A / rifamorpholine B / rifamorpholine C / rifamorpholine D / rifamorpholine E | - | - | - | + (9) | - |
| **NRPS** | isonitrile lipopeptides | +(100) | - | - | - | - |
| **NRPS** | mycobactin | +(100) | - | - | - | - |
| **NRPS** | cyclomarin D | + (8) | + (8) | - | - | - |
| **NRPS** | Orphan | + | + | + | + | + |
| **NRPS** | paenibactin | - | - | + (83) | - | - |
| **NRPS** | capreomycin IA / capreomycin IB / capreomycin IIA / capreomycin IIB | - | - | + (6) | - | - |

| BGC Type | Most similar known cluster type | ASQP_29 | ASQP_78 | ASQP_80 | ASQP_92 | ASQP_98 |
|---|---|---|---|---|---|---|
| **NRPS** | cysteoamide | - | - | + (72) | - | - |
| **NRPS** | scabichelin | - | - | + (90) | - | - |
| **NRPS** | rimosamide | - | - | - | + (42) | - |
| **NRPS** | rhizomide A / rhizomide B / rhizomide C | - | - | - | - | + (100) |
| **NRPS** | istamycin | - | - | - | - | + (4) |
| **NRPS** | thioviridamide | - | - | - | - | + (10) |
| **NRPS** | A40926 | - | - | - | - | + (5) |
| **NRPS,NRPS-like** | vazabitide A | + (4) | - | - | - | - |
| **NRPS,NRPS-like** | teleocidin B1 | - | + (50) | - | - | - |
| **NRPS,NRPS-like** | deimino-antipain | - | - | - | + (66) | - |
| **NRPS,NAPAA** | stenothricin | - | - | - | - | + (13) |
| **NRPS,oligosaccharide ,T2PKS,PKS-like,RRE-containing** | prejadomycin / rabelomycin / gaudimycin C / gaudimycin D / UWM6 / gaudimycin A | - | - | - | + (29) | - |
| **NRPS,RRE-containing,phosphonate,T1PKS,NRPS-like** | fosfazinomycin A | - | - | - | + (89) | - |
| **NRPS,T1PKS** | maklamicin | - | - | - | + (13) | - |
| **NRPS,T1PKS** | diisonitrile antibiotic SF2768 | - | - | - | - | + (66) |
| **NRPS,T1PKS,other** | polyoxypeptin | + (64) | + (64) | - | - | - |
| **NRPS-like** | Orphan | + | - | - | - | - |
| **NRPS-like** | leinamycin | - | - | + (4) | - | - |
| **NRPS-like** | ulleungmycin | - | - | - | + (22) | - |
| **NRPS-like,T1PKS** | spiramycin | - | - | + (6) | - | - |
| **NRPS-like,T1PKS** | WS9326 | - | - | - | - | + (7) |
| **other** | himastatin | + (12) | + (12) | - | - | - |
| **other,terpene** | A-503083 A / A-503083 B / A-503083 E / A-503083 F | - | - | - | + (7) | - |
| **phenazine** | lomofungin | - | - | - | + (39) | - |
| **PKS-like,butyrolactone** | natamycin | + (9) | + (9) | - | - | - |
| **redox-cofactor** | TP-1161 | + (12) | - | - | - | - |
| **redox-cofactor** | lankacidin C | - | - | + (13) | - | + (13) |
| **redox-cofactor** | Orphan | - | - | + | - | - |
| **RiPP-like** | Orphan | + 3 clusters | + 2 clusters | + | +3 clusters | + 2 clusters |
| **RiPP-like** | informatipeptin | + (28) | + (28) | + (42) | - | + (42) |
| **RRE-containing** | A-500359 A / A-500359 B | - | - | - | + (5) | - |
| **siderophore** | desferrioxamin B / desferrioxamine E | + (83) | + (83) | + (83) | + (100) | + (83) |
| **siderophore** | Orphan | + 2 clusters | + 2 clusters | + 2 clusters | + | + |
| **siderophore** | paulomycin | - | - | + (9) | - | - |

| BGC Type | Most similar known cluster type | ASQP_29 | ASQP_78 | ASQP_80 | ASQP_92 | ASQP_98 |
|---|---|---|---|---|---|---|
| **siderophore** | ficellomycin | - | - | - | - | + (5) |
| **T1PKS** | glycopeptidolipid | + (10-20) | - | - | - | - |
| **T1PKS** | foxicins A-D | + (12) | + (12) | - | - | + (21) |
| **T1PKS** | maduropeptin | - | - | + (3) | - | - |
| **T1PKS** | fluostatin | - | - | - | - | + (6) |
| **T1PKS,NRPS-like** | herboxidiene | + (2) | - | - | - | + (4) |
| **T2PKS,hglE-KS,T1PKS** | spore pigment | + (83) | + (83) | - | - | - |
| **T2PKS,oligosaccharide** | saprolmycin E | + (83) | + (83) | - | - | - |
| **T2PKS,other,indole** | spore pigment | - | - | - | - | + (83) |
| **T2PKS** | spore pigment | - | - | + (75) | - | - |
| **T2PKS** | collinomycin | - | - | + (96) | - | - |
| **T2PKS** | kinamycin | - | - | - | - | + (45) |
| **T3PKS** | germicidin | + (100) | + (100) | - | - | - |
| **T3PKS** | alkylresorcinol | + (100) | + (100) | + (100) | + (100) | + (100) |
| **T3PKS** | herboxidiene | - | - | + (8) | - | + (8) |
| **T3PKS** | napyradiomycin A80915C | - | - | + (6) | - | - |
| **T3PKS** | violapyrone B | - | - | - | + (28) | - |
| **T3PKS** | naringenin | - | - | - | + (100) | - |
| **T3PKS,T1PKS** | methylated alkyl-resorcinol / methylated acyl-phloroglucinol | + (100) | - | - | - | - |
| **T3PKS,thioamitides** | BE-7585A | + (11) | - | - | - | - |
| **terpene** | geosmin | + (100) | + (100) | - | + (100) | + (100) |
| **terpene** | hopene | + (15-92) | + (92) | + (92) | + (84) | + (92) |
| **terpene** | avermitilol | + (100) | + (100) | - | - | + (100) |
| **terpene** | albaflavenone | + (100) | + (100) | + (100) | - | + (100) |
| **terpene** | Orphan | - | - | + | - | - |
| **terpene** | cinnamycin | - | - | - | + (9) | - |
| **terpene** | pristinol | - | - | - | + (100) | - |
| **terpene,butyrolactone** | γ-butyrolactone | - | - | + (100) | - | - |
| **terpene,lanthipeptide-class-ii** | 2-methylisoborneol | - | - | - | - | + (100) |
| **terpene,melanin** | melanin | - | - | + (71) | - | - |
| **thioamide-NRP,blactam,NRPS,prodigiosin** | tabtoxin | + (13) | + (13) | - | - | - |
| **thioamitides** | Orphan | + | + | - | - | - |
| **thiopeptide** | mithramycin | + (5) | - | - | - | - |
| **thiopeptide** | actinomycin D | - | + (10) | - | - | - |
| **thiopeptide,RiPP-like,thioamide-NRP,ladderane,NRPS** | radamycin / globimycin | - | - | - | + (77) | - |
